# Cognitive reserves hold back age-related upregulation of neural activities in speech-in-noise perception

**DOI:** 10.1101/2024.05.27.596069

**Authors:** Lei Zhang, Bernhard Ross, Yi Du, Claude Alain

## Abstract

During cognitive tasks, increased frontoparietal neural activity and functional connectivity are commonly observed in older adults. Cognitive reserves accrued from positive life choices can provide additional neural resources to cope with aging. However, how cognitive reserves interact with upregulated neural activity in older adults is poorly understood. We measured brain activity with fMRI during a speech-in-noise task and assessed whether cognitive reserve accumulated from long-term musical training bolsters or holds back age-related upregulated activity. Older musicians exhibited less upregulation of task-induced functional connectivity than older non-musicians in auditory dorsal regions, which predicted better behavioral performance in older musicians. These results suggest that cognitive reserve may hold back neural recruitment. Besides functional connectivity strength, we also found that older musicians showed more youth-like fine spatial patterns of functional connectivity than older non-musicians. Our findings enlightened the intricate interplay between cognitive reserve and age-related upregulated activity during speech in noise perception.

**Significance Statement:** Understanding the interplay between cognitive reserve and age-related neural changes is vital for developing strategies to mitigate cognitive decline in older adults. This study reveals that long-term musical training, which provides cognitive reserve for speech perception, helps older musicians preserve youth-like levels of functional connectivity and fine-scale activity patterns in the auditory dorsal stream during speech-in-noise perception. Unlike older non-musicians, who exhibit age-related neural upregulation, older musicians maintain youth-like connectivity patterns akin to those of younger adults, resulting in superior speech-in-noise perception. These findings support the "Hold-back upregulation" hypothesis and suggest that musical training counteracts age-related declines by preserving youthful brain function. The results offer valuable insights for designing interventions to enhance aging populations’ cognitive function and communication abilities.

## Introduction

Normal aging is associated with sensory and cognitive decline. Current theories posit that older adults may automatically or voluntarily (e.g., spending more effort) engage in strategies to mitigate age-related declines in perception and cognition (1–5). The recruitment of neural activity and strengthening of functional connectivity is thought to index older adults’ compensatory strategy aiming to maintain optimal cognitive performance (6, 7).

According to the Posterior-Anterior Shift in Aging and Compensation-Related Utilization of Neural Circuits model (8, 9), older adults tend to activate frontoparietal brain regions more extensively than younger adults during cognitive tasks (**Fig. 1A**). However, the degree of increased activity in these frontoparietal regions varies significantly among older adults and may be influenced by cognitive and brain reserve (6, 7, 10). The Scaffolding Theory of Aging and Cognition model suggests that positive lifestyle choices, such as musical training, higher levels of education, and bilingualism, contribute to cognitive and brain reserve, which represents the accumulation of cognitive and neural resources before the onset of age-related brain changes (4, 6) (**Fig. 1B, C**). Despite this, how cumulative reserves influenced by positive lifestyle factors impact the recruitment of neural activity in older populations remains controversial. Some studies have found that developing expertise is associated with reduced regional brain activity, which is explained by increased neural efficiency (6, 11, 12). In contrast, others report stronger regional brain activity, attributed to greater functional capacity (13–15). Notably, no studies to date have examined these effects in older adults.

**Fig. 1:**
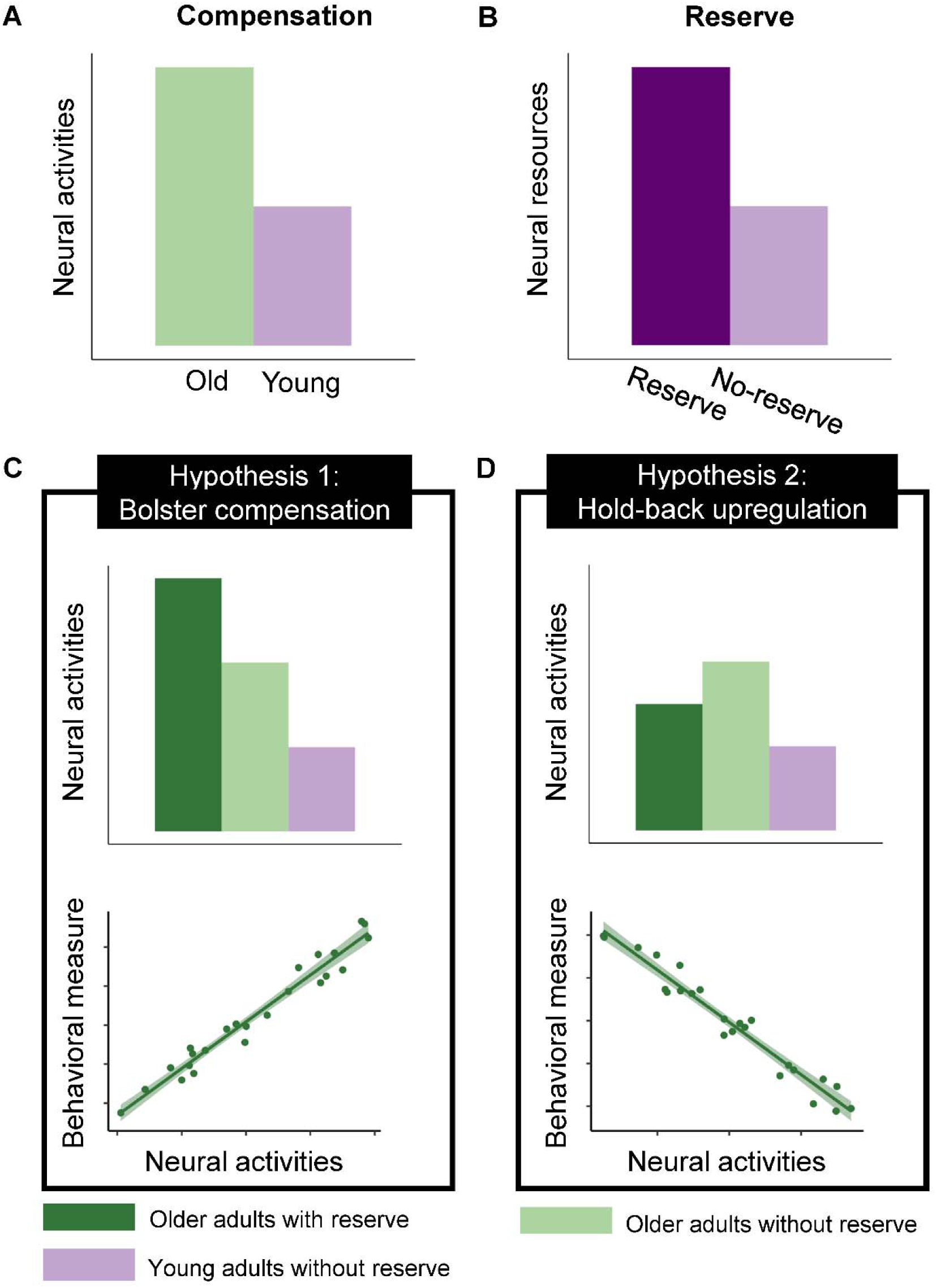
I**l**lustration **of “Bolster compensation” and “Hold-back upregulation” hypotheses**. **A-B**, neural activity that supports age-related neural compensation and neural reserve. **C-D**, Hypothetical change in neural activities and brain-behavior correlation pattern due to the interplay between neural reserve and neural compensation.

We propose two hypotheses regarding how accumulated reserve from long-term musical training interacts with upregulated neural activity when older adults perform speech-in-noise (SIN) perception tasks. The "Bolster Compensation Hypothesis" (**Fig. 1D**) suggests that brain reserve, as a cumulative enhancement of neural resources, strengthens compensatory upregulation of neural activity in older adults during cognitive tasks, with both mechanisms working to increase neural activity and support cognitive performance (6). Conversely, the "Hold-Back Upregulation Hypothesis" (**Fig. 1E**) posits that older adults with greater cognitive reserve display neural activity levels more comparable to those of younger adults, as cognitive reserve provides additional resources that mitigate an age-related decline in perception and cognition (4, 5). The more closely these activity levels resemble those of younger adults, the better the cognitive performance.

In this functional magnetic resonance imaging (fMRI) study, we focused on the complex activity of musical training because it requires sensory-motor integration and provides an ideal model for investigating experience-related plasticity (16–19). Moreover, musicians show cognitive and brain reserve for auditory perception (13, 15, 20–26). Prior research revealed enhanced sensorimotor integration, better speech signal encoding and more robust functional connectivity in the auditory dorsal brain region during speech processing (27–31). Older adults over-recruit dorsal stream brain regions to compensate for deficits in SIN perception (32, 33). Musicians showed greater activity and functional connectivity in dorsal stream regions when they perceive music and SIN (13, 20, 34), which indicates that musical training promotes cognitive reserve in the dorsal stream for SIN perception. These findings suggest that neuroplasticity in the auditory dorsal stream underlies the effects of long-term musical training and aging on SIN perception. Therefore, we will focus on the neural response within the auditory dorsal stream, which includes auditory, inferior parietal, dorsal frontal motor, and frontal motor areas, supporting sound-to-action mapping and sensorimotor integration during speech processing (35, 36).

To test the proposed hypotheses, we invited 25 older musicians (OMs), 25 older non-musicians (ONMs), and 24 young non-musicians (YNMs) to identify syllables in noise (signal-to-noise ratio (SNR): -8, 0, +8 dB). We conducted region of interest (ROI)-based generalized psychophysiological interaction (gPPI) analyses from bilateral auditory seeds (posterior superior frontal gyrus (pSTG)) to bilateral auditory dorsal stream regions (supramarginal gyrus (SMG), supplementary motor area (SMA), superior part of precentral gyrus (PrCGsup), speech motor areas (SM) including inferior part of precentral gyrus (PrCGinf) and opercular part of inferior frontal gyrus (IFGop)). We also analyzed the resting-state functional connectivity (RSFC) of the three groups to investigate whether the hypotheses are supported solely by task-induced activity or intrinsic activity observed during the resting state. Additionally, we extracted blood-oxygen-level-dependent (BOLD) activation patterns in these ROIs, as most previous studies recognized neural compensation and neural reserve based on regional activations (6, 7, 32, 33).

Observing upregulated neural activity in ONMs compared to OMs supports the “Hold-back upregulation hypothesis.” In this case, we expect the level of neural activities in OMs to range between ONMs and YNMs. We anticipate the difference between OMs and YNMs to be smaller than between ONMs and YNMs. We also expect a negative correlation between neural activities in OMs and their behavioral performance. However, an extra upregulation of the neural activities in OMs would support the “Bolster compensation hypothesis.” This would be accompanied by a larger difference between OMs and YNMs than the difference between ONMs and YNMs and a positive correlation between neural activities in OMs and their behavioral performance.

## Results

### Musical training mitigated age-related declines in SIN perception

First, we analyzed the accuracy of syllable identification to examine whether long-term musical training protected against age-related declines in SIN perception among older adults. We performed two-way mixed-design analyses of variance (ANOVA).

We found a significant group effect, an SNR effect, and an interaction between group and SNR (group: F(2,71) = 28.65, P < 0.001; SNR: F(2,71) = 63.78, P < 0.001; Interaction: F(4,71) = 4.24, *P* = 0.004). OMs showed better behavioral performance than ONMs under two higher SNR conditions (SNR 8: t(71) = 3.89, P_fdr_ < 0.001; SNR 0: t(71) = 3.29, P_fdr_ = 0.002; SNR -8: t(71) = -0.26, P_fdr_ = 0.794), albeit worse than YNMs (SNR 8: t(71) = -3.15, P_fdr_ = 0.002; SNR 0: t(71) = -2.88, P_fdr_ = 0.005; SNR -8: t(71) = -5.59, P_fdr_ < 0.001), and YNMs performed better than ONMs (SNR 8: t(71) = 7.00, P_fdr_ < 0.001; SNR 0: t(71) = 6.13, P_fdr_ < 0.001; SNR -8: t(71) = 5.33, P_fdr_ < 0.001) (**Fig. 2**). Although the SNR effect was significant, the behavioral performance differences among older adults, particularly non-musicians, across the three SNR levels were minimal (ONMs: 35%-43%; OMs: 34%-52%). We focus solely on the group effect in the following analyses.

**Fig. 2:**
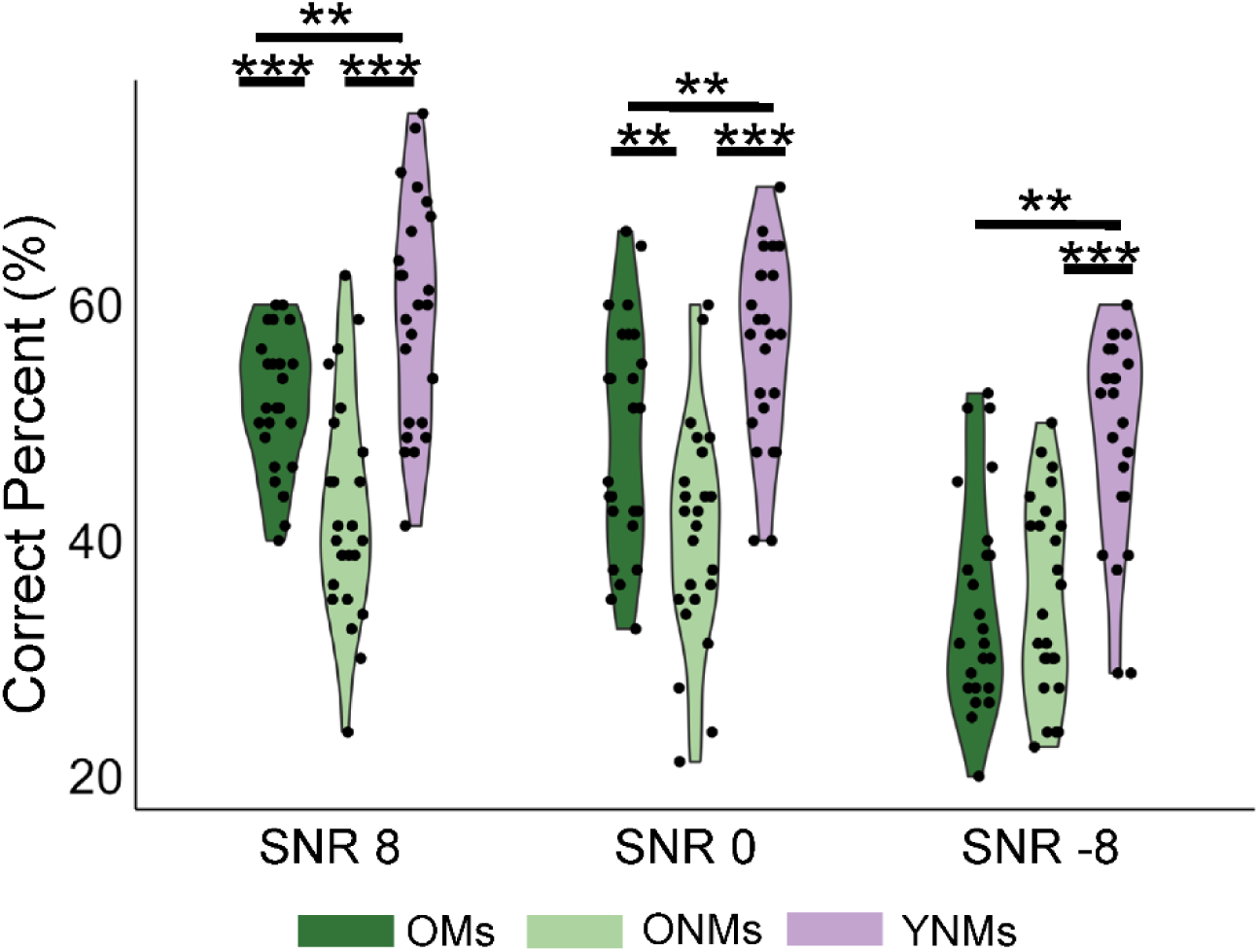
B**e**havioral **performance of the three groups under three SNR conditions**. Violin plots and individual data points of behavioral performance. OMs performed worse than YNMs under all SNRs, but better than ONMs under SNR 8 and SNR 0. ***P_fdr_ < 0.001, **P_fdr_ < 0.01. OMs, older musicians; ONMs, older non-musicians; YNMs, young non-musicians.

### ONMs showed upregulated task-induced functional connectivity (TiFC) during SIN perception

We performed gPPI analysis to investigate how aging and musical training modulated the TiFC within the bilateral auditory dorsal streams during SIN perception. According to the dual model of auditory processing (35–37) and previous results of speech perception studies (13, 31, 35, 38), we used bilateral posterior STG as seeds in the auditory dorsal streams. Target ROIs included bilateral SMG, SMA, PrCGsup, and speech motor areas composed of PrCGinf and IFGop (**Fig. 3A, B**). Two-way mixed-design ANOVAs (group * SNR) were performed to investigate the group effect on TiFC in the bilateral auditory dorsal streams. Outliers, defined as data points beyond two standard deviations, were replaced with the group average.

**Fig. 3:**
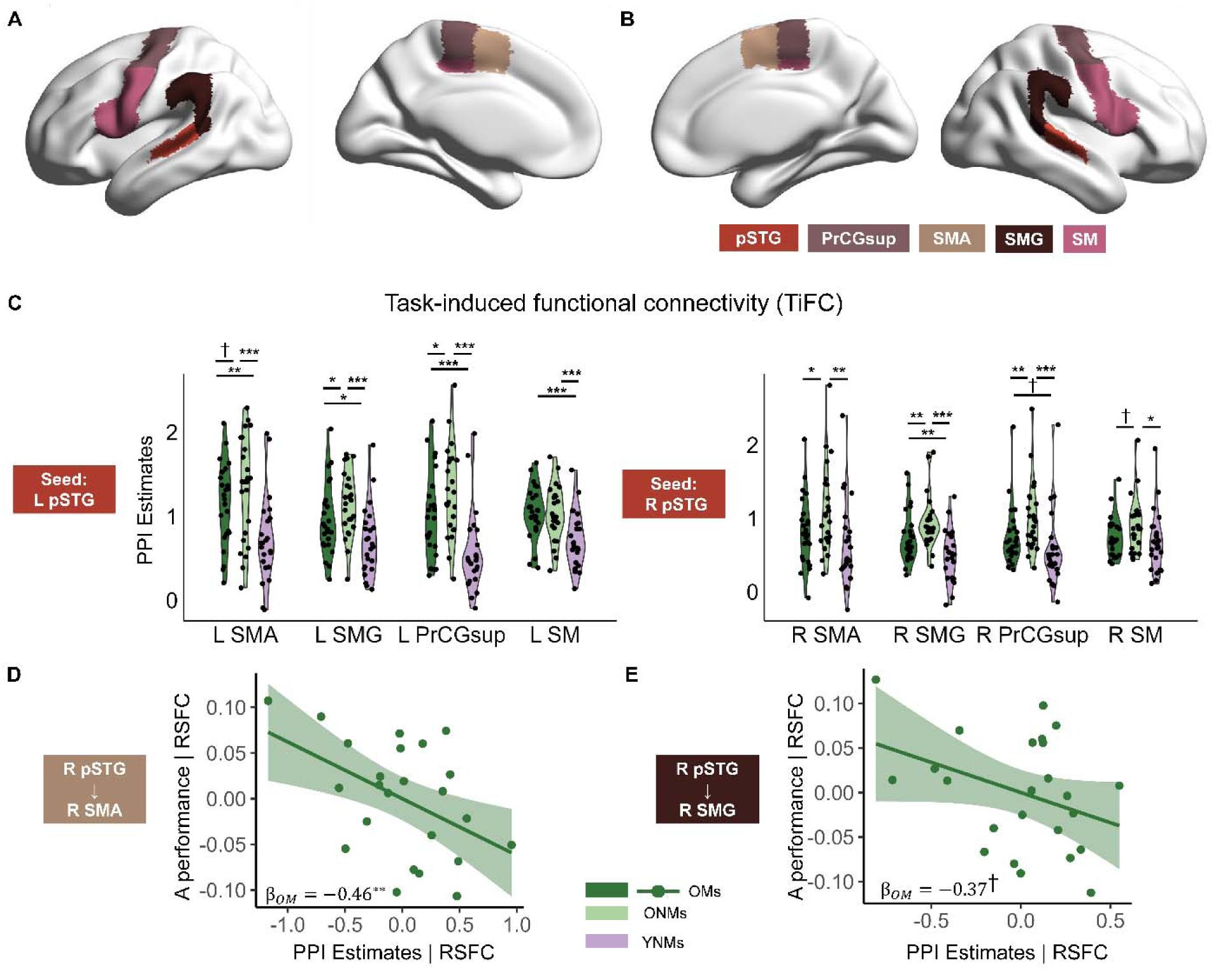
T**i**FC **of the three groups, and brain-behavior correlations. A**, left and **B**, right ROIs used in PPI analysis. Bilateral posterior superior temporal gyri (pSTG) were the seed ROI. Bilateral auditory dorsal stream regions, including supramarginal gyrus (SMG), supplementary motor area (SMA), superior part of precentral gyrus (PrCGsup), and speech motor areas (SM) were the target regions. **C**, ONMs showed upregulated task-induced functional connectivity (TiFC) in bilateral dorsal streams compared to YNMs, while OMs exhibited TiFC resembling YNMs. Violin and point plots show the individual TiFC in specific target regions. Error bars indicate the SEM. **D-E**, TiFC strength of OMs was negatively correlated to the behavioral performance after controlling for resting-state functional connectivity (RSFC). ***P_fdr_ < 0.001, **P_fdr_ < 0.01, *P_fdr_ < 0.05, †P_fdr_ < 0.1; OMs, older musicians; ONMs, older non-musicians; YNMs, young non-musicians; L, left; R, right.

We found a main effect of group on TiFC within the bilateral auditory dorsal streams (LpSTG-LSMA, LSMG, LPrCGsup, LSM; RpSTG-RSMA, RSMG, RPrCGsup, RSM). For connectivity that showed a significant group effect after FDR correction, pairwise comparisons revealed stronger TiFC in ONMs than young adults (LpSTG-LSMA, LSMG, LPrCGsup, LSM; RpSTG-RSMA, RSMG, RPrCGsup, RSM, **Fig. 3C**, **Supplementary Table 1**).

Additionally, we employed robust linear mixed-effects analysis to investigate the effect of group on the left and right hemisphere TiFC after controlling for participants and brain regions. We found stronger overall TiFC in ONMs than YNMs (left: β_group_ = 0.56, *P* < 0.001; right: β_group_ = 0.45, *P* < 0.001).

### Connectivity strength in OM was more akin to YNM than ONM

Both OMs and ONMs showed stronger connectivity than YNMs in the left hemisphere (LpSTG-LSMA, LSMG, LPrCGsup, LSM, **Fig. 3C**, for statistics, see **Supplementary Table 1**). However, OMs showed more youth-like TiFC than ONMs in certain ROIs (LpSTG-LSMG, LPrCGsup). Using a nonparametric bootstrap approach with 10,000 iterations, we found that the difference between OMs and YNMs across the left ROIs was significantly smaller than the difference between ONMs and YNMs (95% Bootstrap CI: 0.59 1.21).

ONMs showed stronger connectivity in the right hemisphere than OMs (RpSTG-RSMA) and YNMs (RpSTG-LSMA, RPrCGsup, RSMG), but differences between OMs and YNMs were not significant except for the connectivity between RpSTG and RSMG. We also found that the difference between OMs and YNMs across the right ROIs was significantly smaller than the difference between ONMs and YNMs (95% Bootstrap CI: 0.35 0.95). Additionally, with a linear mixed model, regardless of brain regions, we found stronger TiFC in ONMs than OMs (β_group_ = 0.21 *P* = 0.023) in the right hemisphere. OMs also showed stronger TiFC than YNMs (β_group_ = 0.23 *P* = 0.021).

Moreover, we observed a negative correlation between TiFC strength and behavioral performance in OMs in the right SMA (β_OMs_ = -0.46, *P* = 0.008, **Fig. 3D**) and a marginally negative correlation in the right SMG (β_OMs_ = -0.37, *P* = 0.078, **Fig. 3D**), when resting-state functional connectivity (RSFC) was controlled as a covariate. In the right PrCGsup, the TiFC-behavior correlation was also negative but not significant (β_OMs_ = -0.32, *P* = 0.163). No other correlations were found. That is, the connectivity strengths of OMs who showed better behavioral performance were more akin to those of young adults, especially in the right hemisphere. Thus, long-term musical training provides the cognitive reserve to mitigate age-related declines in SIN perception. The TiFC strength supports the “Hold-back upregulation” hypothesis.

### OMs showed higher spatial alignment of TiFC to YNMs than ONMs

We performed inter-subject spatial correlation (ISPC) to further explore whether OMs also showed more youth-like TiFC in fine spatial patterns in the regions exhibiting significant or marginally significant lower TiFC strength than ONMs. ISPC measures the spatial neural alignment of multivoxel TiFC spatial patterns of the older brain to that observed in younger brains (**Fig. 4A**). We found that OMs showed higher ISPC to YNMs than ONMs in left PrCGsup (t(46) = 2.74, P_fdr_ = 0.044, **Fig. 4B**). That is, OMs showed both more youth-like TiFC strength and spatial TiFC pattern than ONMs. More importantly, higher ISPC of OMs to YNMs negatively correlated with the TiFC strength after controlling for RSFC (β_OMs_ = -0.37, *P* = 0.029, **Fig. 4C**). This suggests that the more youth-like spatial pattern is related to greater “Hold-back upregulation.”

**Fig. 4:**
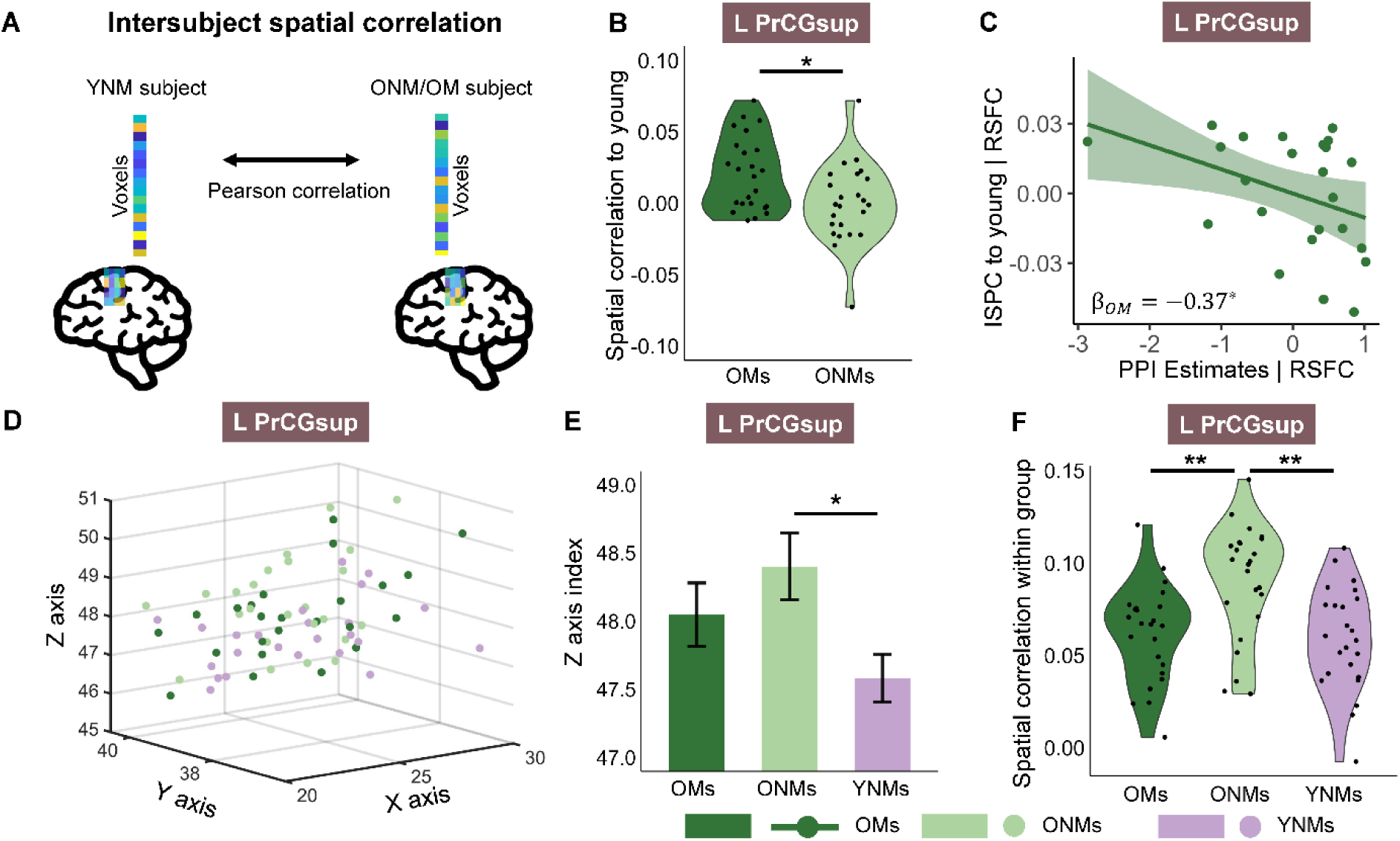
S**p**atial **alignment of TiFC to YNMs and within group**. **A**, Illustration of the calculation of intersubject spatial correlation; see Methods for details. **B-C**, OMs showed greater spatial alignment to YNMs than ONMs in left PrCGsup (**B**), and greater spatial alignment to YNMs predicted lower TiFC strength in OMs (**C**). **D-E**, Mean 3D coordinates of the voxels that showed top 10% TiFC were plotted in **D**. Voxels of ONMs exhibited significantly different positions in Z axis compared to YNMs, while no difference was found between OMs and YNMs (**E**). **F**, ONMs showed greater spatial correlation within the group than OMs and YNMs. Bar plots show the group mean of specific Z-axis index in L PrCGsup. Error bars indicate the SEM. *P_fdr_ < 0.05, **P_fdr_ < 0.01; OMs, older musicians; ONMs, older non-musicians; YNMs, young non-musicians; L PrCGsup, left superior part of precentral gyrus.

Next, we explored how the TiFC spatial pattern in left PrCGsup was shifted in ONMs. By extracting the mean coordinate of the top ten percent voxels, we could examine whether the voxel positions were different between the two older adult groups and YNMs (**Fig. 4D**). We found a significant difference between ONMs and YNMs in the Z-axis direction (t(46) = 2.73, P_fdr_ = 0.027, **Fig. 4E**). Since the Z-axis increases from inferior to superior, the top voxels positions in ONMs were superior to those in YNMs. As expected, we found no difference between OMs and YNMs (t(46) = 1.60, P_fdr_ = 0.175, **Fig. 4E**).

We also calculated ISPC within groups. A higher within-group ISPC denotes that the TiFC spatial patterns are more similar within this group. We found that ONMs exhibited a higher ISPC than YNMs and OMs in left PrCGsup (YNM: t(46) = 3.71, P_fdr_ = 0.002; OM: t(46) = 3.46, P_fdr_ = 0.003, **Fig. 4F**). However, no difference was found between OMs and YNMs (t(46) = 0.37, *P* = 0.715, **Fig. 4F**). In summary, ONMs’ TiFC spatial pattern consistently deviated more from YNMs than OMs.

### Older adults showed stronger intrinsic RSFC in the auditory dorsal stream than young adults

We further examined whether the two hypotheses were supported by intrinsic functional connectivity itself by extracting RSFC in auditory dorsal stream regions seeded from bilateral pSTG. As illustrated in **Fig. 5**, OMs demonstrated stronger RSFC than YNMs (LpSTG-LSMA, LSMG, LprCGsup; RpSTG-RSMA, RSMG, RPrCGsup, RSM, for statistics, see **Supplementary Table 2**), with no significant difference observed between OMs and ONMs. ONMs also showed stronger RSFC than YNMs (LpSTG-LPrCGsup; RpSTG-RPrCGsup, for statistics, see **Supplementary Table** 2).

**Fig. 5:**
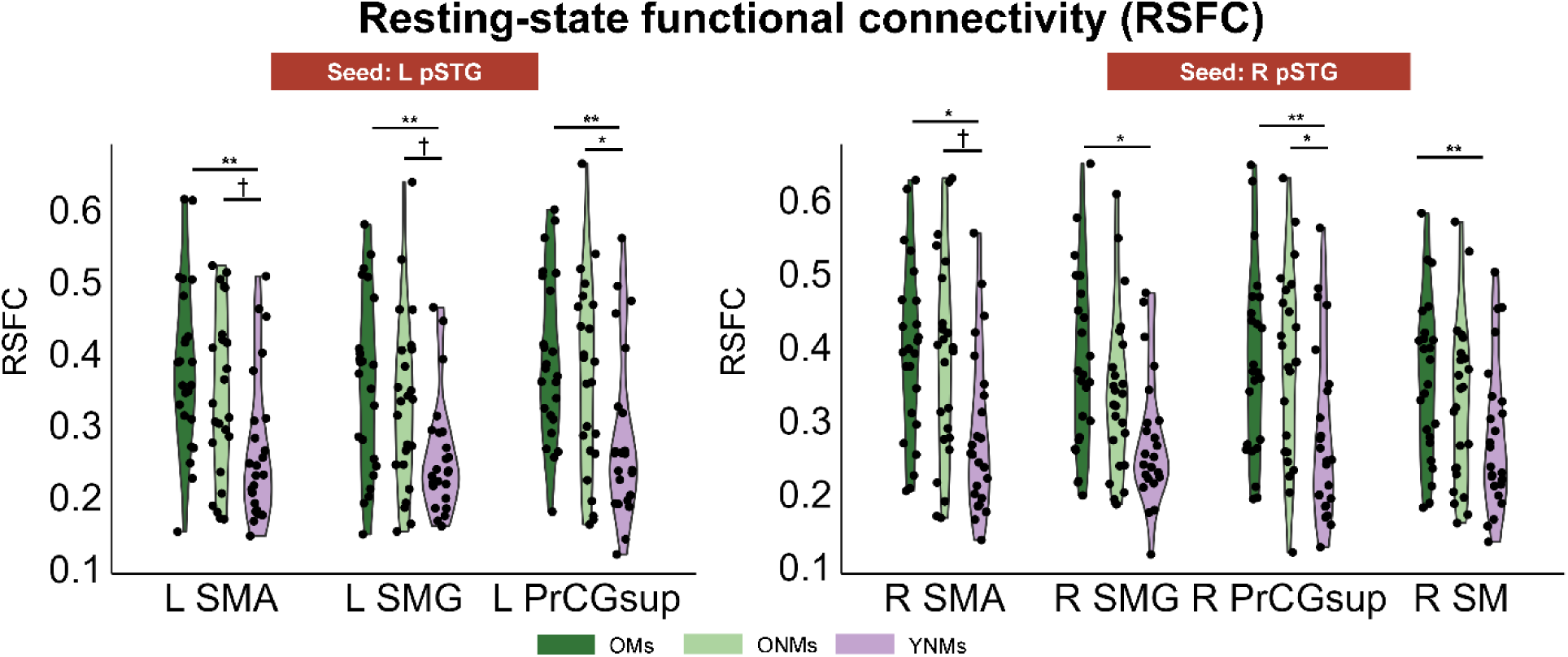
B**i**lateral **RSFC of the three groups.** ONMs and OMs showed upregulated RSFC in bilateral dorsal streams compared to YNMs. Violin and point plots show the individual RSFC in specific target regions. Error bars indicate the SEM. **P_fdr_ < 0.01, *P_fdr_ < 0.05, †P_fdr_ < 0.1, OMs, older musicians; ONMs, older non-musicians; YNMs, young non-musicians; pSTG, posterior part of superior temporal gyrus; SMG, supramarginal gyrus; SMA, supplementary areas; SM, speech motor areas; PrCGsup, superior part of precentral gyrus; L, left; R, right.

However, we did not find any significant RSFC-behavior correlation in both OMs and ONMs groups. Therefore, age-related changes in intrinsic functional connectivity within the dorsal steam, as measured with RSFC, has little impact on task performance.

### No group difference in BOLD activation was found

We extracted BOLD activation in auditory dorsal stream regions to assess the two hypotheses regarding regional activation levels. However, we did not observe any main effect of BOLD activation difference between the three groups (for statistics, see **Supplementary Table 3**).

## Discussion

This study examined how cognitive reserve accumulated through long-term musical training interacts with age-related neural compensation during SIN perception. We found reduced age-related declines in SIN performance among OMs. As expected, ONMs showed the typical age-related compensatory upregulation of TiFC in bilateral auditory dorsal streams. Notably, OMs exhibited a connectivity pattern in the bilateral auditory dorsal stream that resembled YNMs, with connectivity strength in the right dorsal stream correlating with SIN perception.These findings support our “Hold-back upregulation” hypothesis, where cognitive reserve from musical training promotes a more youthful functional connectivity pattern, leading to superior behavioral outcomes. By further investigating fine spatial alignment of TiFC, we found that OMs exhibited more youth-like spatial pattern of TiFC, particularly between LpSTG and LPrCGsup, whereas ONMs showed a more consistently deviating TiFC spatial pattern alignment to YNMs compared to OMs. Lastly, we found increased intrinsic RSFC in older adults but no significant brain-behavior correlation with RSFC. Overall, our findings shed light on the intricate interplay between age-related upregulated neural activity and cognitive reserve. Reserve accrued from long-term musical training holds back the age-related upregulation of neural activities both in terms of strength level and fine spatial pattern such that OMs present youth-like listening skills.

We found a decline in SIN perception associated with aging, consistent with previous studies (20, 39). Previous studies have highlighted the upregulation in frontal BOLD activity as a compensatory mechanism during SIN perception in ONMs (32, 33). However, conflicting findings have been reported in other studies (40–43). In the current study, ONMs did not show increased regional BOLD activation during SIN perception. Instead, they exhibited heightened TiFC strength within bilateral auditory dorsal streams. The interpretation of upregulated neural activity varies depending on brain-behavior correlations. A positive correlation suggests neural compensation, whereas a negative correlation indicates neural dysfunction (6, 7). However, in this study, we did not find a consistent positive or negative relationship between higher connectivity strength and better behavioral performance in ONMs. Therefore, it remains to be determined whether the upregulation of functional connectivity within auditory dorsal streams in ONMs indicates a compensatory strategy.

By investigating the neural responses of OMs, we showed how accumulated cognitive reserve from long-term musical training interacts with age-related neural upregulation. Previous studies in young adults have found that musicians show greater TiFC from auditory to motor areas during SIN perception, supporting their superior SIN perception performance (13). Additionally, musicians displayed enhanced sensorimotor integration due to intensive sensory-motor interaction during musical training (15, 16, 18, 44). In the bilateral hemisphere, we observed that OMs showed a connectivity profile more akin to YNMs than ONMs, based on the mean TiFC strength in most regions and the group difference across regions. In addition, lower connectivity strength in right brain predicts better behavioral performance in OMs. These results support the “Hold-back upregulation” hypothesis in the right brain, suggesting that the cognitive reserve provided by long-term musical training preserves a youth-like brain connectivity strength, which contribute to better behavioral performance. This is consistent with our previous findings using the same dataset, which show that OMs maintain better SIN perception performance by preserving youth-like representations in sensorimotor regions (20). Previous studies have found that young musicians exhibit both functional and structural advantages of the right dorsal stream compared to YNMs, which may explain the potentially greater “Hold-back upregulation” in the right hemisphere in OMs (13, 45–47).

This study significantly extends prior research (20) by showing both averaged TiFC strength and fine spatial pattern of TiFC during SIN processing, collectively suggesting that cognitive reserve from long-term musical training mitigates age-related declines in SIN perception by preserving youth-like TiFC patterns within task-related neural networks. Previous studies on age-related changes in brain function have compared univariate activities and multivoxel representations of presented stimuli (6, 7, 32). By correlating the spatial patterns between older and young adults, referred to as intersubject spatial correlation, we directly examine the similarity of fine activation patterns between age groups. Both prior and current studies have found that youth-like neural activity patterns are associated with preserved behavioral performance and neural activity levels in older adults (20). We believe that quantifying the similarity of fine activity patterns between young and older brains provides a valid and intuitive index of functional aging.

According to the dual-stream model of speech perception, the auditory dorsal stream supports sound-to-action mapping during speech processing (35, 36). Previous studies have shown that task-related neural activities increase in auditory dorsal stream regions, including the frontal motor areas (e.g., premotor cortex and IFG), to compensate for deteriorating auditory signals via sensorimotor integration(31, 32, 48). Deteriorated auditory function due to aging can also be regarded as an adverse condition (49). Learning to play a musical instrument is a long-term process that requires intensive sensorimotor integration among auditory, visual, and motor system. Musicians have been shown to have an advantage in auditory-motor integration during SIN perception (13, 20). Therefore, the auditory dorsal stream is central to both age-related upregulation and the neural reserve from musical training, making it ideal for investigating how these two mechanisms interact. To our knowledge, this study is the first to directly investigate how accumulated cognitive reserve influences age-related upregulated neural activity. Future studies should further test this hypothesis using different cognitive tasks, such as memory and attention tasks, and investigate other sources of reserve, such as physical exercise and bilingualism. Additionally, examining how different types of learning transfer may influence such interactions will be important for generalizing the findings.

RSFC is an excellent approach for examining intrinsic connectivity of participants independent of specific tasks. Previous studies have found that musicians showed more robust RSFC than non-musicians in speech-related brain networks (50–52). We also examined the intrinsic functional connectivity in the three groups and included RSFC as a covariate in the brain-behavior correlation analysis. We found that both ONMs and OMs showed stronger RSFC compared to YNMs. Previous research has found decreased within-network connectivity but increased between-network connectivity in older adults compared to young adults (for reviews, see (53)). This aligns with our findings that older adults’ connectivity between auditory and frontal motor areas increased. However, whether RSFC is necessarily correlated with behavioral performance remains unclear (51, 54). In this study, since we did not find any significant RSFC-behavior correlation in either older groups, it cannot be concluded that the stronger intrinsic functional connectivity observed in older adults serves as a compensatory mechanism.

Further studies are need to explore the relationship between RSFC and SIN perception in older adults.

In conclusion, this fMRI study offers new insights into the intricate interplay between neural reserve and age-related upregulated activity in SIN perception among older adults. Specifically, it examines the newly proposed “Hold-back upregulation” and “Bolster compensation” hypotheses within this context. Our findings suggest that long-term musical training mitigates age-related decline in SIN perception by enhancing cognitive reserve, which interacts with upregulation of tasked-induced functional connectivity in ways consistent with the “Hold-back upregulation” hypothesis. Our results underscore the importance of considering both cognitive reserve and age-related upregulated neural activity in understanding cognitive performance, like SIN perception, in older adults. These findings may inform interventions aimed at preserving cognitive function and improve communication outcomes in aging populations.

## Methods

### Participants

Twenty-five OMs, 25 ONMs, and 24 YNMs participated in this study. One OM was excluded from the fMRI analysis because of excessive head movements, and another ONM because of left-handedness. All remaining participants were healthy, right-handed, native Mandarin speakers with no history of neurological disorder and normal hearing (average pure tone threshold < 20 dB hearing level between 250 Hz and 4000 Hz) in both ears. All older adults passed the Beijing version of the Montreal Cognitive Assessment (MoCA, ≥26 scores)(55). All OMs started musical training before the age of 23 years (10.90 ± 4.56 years old) and had at least 32 years of musical training (50.88 ± 8.75 years). Also, all OMs practiced regularly in the last three years (12.70 ± 8.99 hours per week). All non-musicians reported less than two years of musical expertise experience. The demographic information of the three groups is summarized in **Table 1**. OMs and ONMs were not different in age, pure tone average (PTA), and MoCA score. However, self-reported education years differed between groups (OMs > ONMs, t = 4.15, *P* < 0.001). We conducted supplementary analyses to control for the effect of years of education on our results (see Supplementary Text of the **Supplementary Materials**). All participants provided written consent before taking part in the study, which was approved by the ethics committee of the Institute of Psychology, Chinese Academy of Science.

**Table 1.**
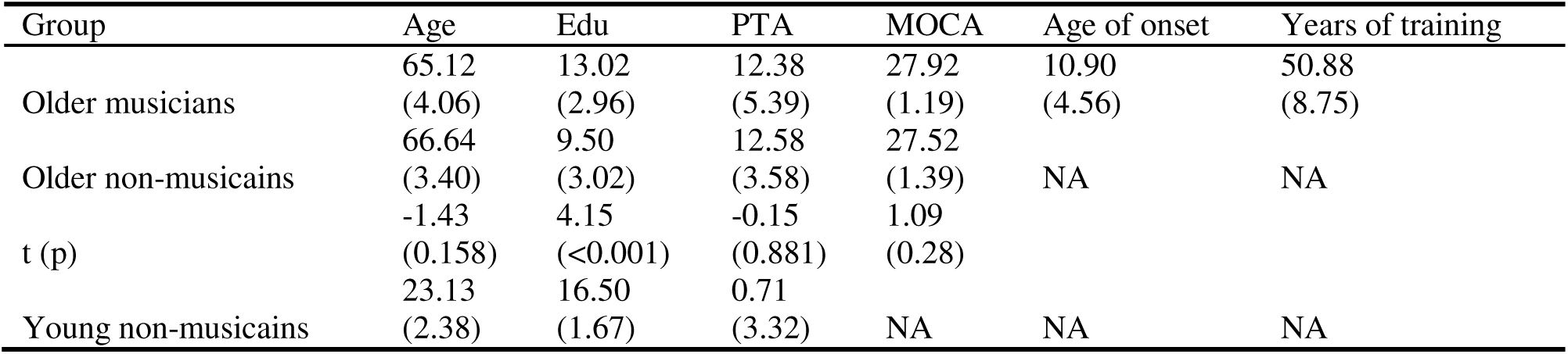
The Group Mean (Standard Deviation) Values and Statistics of Age, Education, Pure Tone Average (PTA) at 250–4000 Hz, MOCA Score, Age of Training Onset, and Years of Music Training in Each Group.

### Stimuli and procedure

The stimuli comprised four naturally pronounced consonant-vowel syllables (/ba/, /da/, /pa/, /ta/) uttered by a 23 years old Chinese female. The utterances were recorded in a soundproof room. The syllable stimuli were about 400 ms in duration, low-pass filtered (4 kHz), and matched for average root-mean-square sound pressure level (for more details, see(20, 31)). The masker was a speech spectrum-shaped noise (4-kHz low-pass, 10-ms rise-decay envelope) representative of 113 sentences by 50 young Chinese female speakers (age range 20 to 26 years). The stimuli were delivered at a level of 90 dB sound pressure level (SPL). The SPL of the masking sounds was modified to create three signal-to-noise ratios (SNRs) of -8, 0, and 8 dB. The stimuli were transmitted using MRI-compatible insert earphones (S14, Sensimetrics Corporation), which attenuated the scanner noise by up to 40 dB.

In the fMRI scanner, participants listened to the speech signals and identified the syllables by pressing the corresponding button using their right-hand fingers (index to little fingers in response to ba, da, pa, and ta in half of the participants or pa, ta, ba and da in the other half of the participants sequentially). Each participant completed four blocks. Each block contained 60 stimuli (20 trials × 3 SNRs), pseudo-randomly presented with an average inter-stimuli-interval of 5 s (4–6 s, 0.5 s step).

### Behavior data analysis

Mixed-design ANOVAs were performed to investigate the main effect of group on behavioral performance. Greenhouse–Geisser correction was applied if the sphericity assumption was violated. FDR correction accounted for multiple comparisons in post hoc analysis. Statistical analyses were performed with the package bruceR (56) in R.

### Imaging data acquisition and preprocessing

Functional imaging data was collected by a 3T MRI system (Siemens Magnetom Trio), and T1 weighted images were acquired using the MPRAGE sequence (TR = 2200 ms, TE = 3.49 ms, FOV = 256 mm, voxel size = 1×1×1 mm). Blood oxygen level-dependent (BOLD) images were acquired using the multiband-accelerated EPI sequence (acceleration factor = 4, TR = 640 ms, TE = 30 ms, slices = 40, FOV = 192, voxel sizes = 3×3×3 mm).

Task-state functional imaging data were preprocessed using AFNI software (57). The first eight volumes were removed. The following preprocessing steps included slice timing, motion correction, aligning functional images with anatomy, spatial normalization (MNI152 space), spatial smoothing with 6-mm FWHM isotropic Gaussian kernel, and scaling each voxel time series to have a mean of 100.

### Univariate general linear model (GLM) analysis

Single-subject multiple-regression modeling was performed using the AFNI program 3dDeconvolve. Six conditions of four syllables and six regressors corresponding to motion parameters were entered into the analysis. TRs would be censored if the motion derivatives exceeded 0.3. For each SNR, the four syllables were grouped and contrasted against the baseline, and BOLD activation was averaged across SNRs.

### Task-induced functional connectivity

gPPI analysis was performed to investigate the TiFC between auditory seeds and other regions in the auditory dorsal stream(58). Auditory seeds were defined as bilateral posterior division of STG labels from the Harvard-Oxford atlas. Target regions were auditory dorsal stream regions, including bilateral SMG, SMA, PrCGsup, and SM area, which comprises preCGinf and IFGop (31, 35, 38). Note that since inter-hemispheric connectivity was not of interest in this study, only intra-hemispheric functional connectivity was discussed in this study (right seed to right target; left seed to left target). Subject-level gPPI analyses were conducted separately for left and right auditory seeds using the AFNI program 3dDeconvolve. PPI regressors, the seed time series and the regressors of the original GLM model were included in the model.

### Resting-state functional connectivity

We also investigated intrinsic functional connectivity by analyzing resting-state functional data. Resting-state data were preprocessed using the GRETNA toolbox (59). After removing the first ten volumes, the remaining 740 volumes entered the following preprocessing steps: slice timing, realignment, spatial normalization, and detrending. Lastly, the normalized images were temporally filtered between 0.02 and 0.1 Hz, and covariates, including cerebrospinal fluid signal, white matter region signal, and head motion parameters, were regressed out.

Pearson correlation coefficients were calculated between voxel-wise time series in auditory dorsal stream regions, the same target ROIs used in gPPI analysis and the mean time series in bilateral auditory seeds to obtain the intrinsic functional connectivity in the auditory dorsal stream.

### ROI-based group analysis

To specifically investigate how aging and musical training modulate regional activities in the auditory dorsal stream during audiovisual speech perception, we performed the ROI analysis in BOLD activation, TiFC and RSFC. Voxel-wise TiFC, RSFC, and BOLD activation were extracted for each ROI. Then, voxels with the top 40% activation or connectivity levels across conditions were selected and averaged for the following statistical analysis(60). Mixed-design ANOVAs (group*SNRs) were performed for A and visual enhancement. FDR correction was applied for multiple comparisons across ROIs and post hoc analyses. Outliers, defined as data points beyond two standard deviations, were replaced with the group average. Correlation analysis was also performed to test brain-behavior correlation.

To determine whether the difference between YNMs and ONMs (/J_ONM-YNM_) was larger/smaller than the difference between YNMs and OMs (/J_OM-YNM_), we compared the two mean differences over left or right ROIs directly and employed a nonparametric bootstrap approach with 10,000 iterations to quantify the uncertainty in this comparison. In each bootstrap iteration, we resampled the data for each group with replacement, preserving the original sample size. For each resampled dataset, we recalculated /J_ONM-YNM_ and /J_OM-YNM_ . From the resulting empirical bootstrap distribution, we derived the 95% confidence interval (CI) by taking the 2.5th and 97.5th percentiles. If this CI did not include zero, it indicated that the difference between YNMs and ONMs was statistically significantly larger/smaller than the difference between OMs and ONMs.

A robust linear mixed model analysis was further performed to investigate the group effect (e.g., ONMs vs. YNMs) on neural indices regardless of brain regions, which showed significant group effects in each hemisphere. The formula was established as follows:

Neural index = group + (1|subject) + (1|brain regions)

The above analyses were performed using the packages “bruceR “ and “robustlmm” in R (56, 61).

### Inter-subject spatial correlation (ISPC) analysis

The TiFC of each voxel within the ROIs that showed significant group difference between OMs and ONMs under each condition was extracted for each participant. Then, we correlated the TiFC of each OM and ONM to that of each YNM for each ROI. The averaged correlation coefficients of each older adult to all YNMs represented the spatial alignment of the young. A higher spatial alignment measure denotes that the subject showed a higher TiFC spatial similarity as young adults. FDR correction was performed for multiple comparisons.

ISPC was also performed within the groups. That is, we correlated TiFC of each OM to that of other OMs for the ROI. The same procedure was also performed for the other two groups. The average correlation coefficients of each participant were calculated and compared to those of other participants within the group. Higher within-group ISPC denotes that the neural activities of the subjects in that group are more homogeneous with each other, which probably suggests that subjects in that group employ similar processing strategies.

### Activity spatial position analysis

Following ISPC, we performed the activity spatial position analysis within the specific ROI to examine whether older adults showed position differences of voxels that showed top TiFC. We extracted voxels that showed the top 10% TiFC strength of each participant within the ROI that showed significant ISPC group differences between OMs and ONMs for a more specific and targeted location analysis. Then, we averaged the x (left to right), y (posterior to anterior), and z (inferior to superior) axis coordinates of the voxels for each participant. T-tests were performed to compare the mean coordinates between groups. FDR correction was performed for multiple comparisons.

## Acknowledgments

We thank Y. Chen and Y. Wu for helping with data collection and X. Jiang for helping with stimuli generation.

## Author contributions

Conceptualization: L.Z., C.A. and Y.D. Methodology: L.Z., and C.A. Investigation: L.Z. Visualization: L.Z., B.R., and C.A. Supervision: Y.D., and C.A., Writing (original draft): L.Z. and C.A. Writing (review and editing): L.Z., B.R., C.A., and Y.D

## Data and materials availability

All data needed to evaluate the conclusions in the paper are present in the paper and/or the Supplementary Materials. The behavioral and fMRI data that support the findings of this study will be available in OSF: https://osf.io/89hbn/.

## Supplementary Materials

### Supplementary Text

#### Effect of years of education on the behavioral performance and neural responses in the audiovisual speech-in-noise perception task

We found that years of education including both formal and informal education were significantly different between ONM and OM. Due to historical reasons in China, all recruited older adults discontinued formal education at the same time (1960s-1970s). Therefore, the self-reported years of education cannot represent the education level of the older subjects reliably.

The years of education were highly correlated with the group variable between OM and ONM (point-biserial correlation r = 0.64), and were correlated with the group variable between older subjects and young subjects (ONM vs. YNM: r = 1; OM vs. YNM: r = 0.74). Therefore, it is not suitable to put the years of education variable into the analyses as a control variable because of the collinearity problem.

To exclude the effect of years of education on our results, we performed supplementary analysis to investigate the effect of years of education on the behavioral performance, BOLD activation, task-induced functional connectivity, and resting-state functional connectivity. We included the years of education in the mixed-designed ANOVA. We found years of education is not significant in all of the analysis (behavioral: all p > 0.522; task-induced functional connectivity: all P > 0.209; resting-state functional connectivity: all P > 0.649). Therefore, years of education did not influence the behavioral performance and neural responses in this study.

---

**Supplementary Table 1.** Mixed-design ANOVA and post hoc analysis of task-induced functional connectivity. The F values and associated P values represent the main effect of the group from a Mixed-design ANOVA. The reported P values have been corrected for multiple comparisons across the ROIs using the FDR method. The t values and associated P values are from post hoc pairwise comparisons with FDR correction.

| Seed: **LSTG** | Target regions | Group main effect<br>$F_{2,69}(P_{fdr})$ | OMs vs. ONMs<br>$t_{69}(P_{fdr})$ | OMs vs. YNMs<br>$t_{69}(P_{fdr})$ | ONMs vs. YNMs<br>$t_{69}(P_{fdr})$ |
| --- | --- | --- | --- | --- | --- |
|  | L SMA | 14.38(<0.001) | 1.68(0.097) | -3.57(0.001) | -5.25(<0.001) |
|  | L SMG | 13.39(<0.001) | 2.55(0.013) | -2.63(0.013) | -5.18(<0.001) |
|  | L PrCGsup | 23.32(<0.001) | 2.58(0.012) | -4.18(<0.001) | -6.77(<0.001) |
|  | L SM | 12.00(0.000) | 0.60(0.553) | -3.91(<0.001) | -4.51(<0.001) |
| Seed: **RSTG** | Target regions |  |  |  |  |
|  | R SMA | 7.60(0.001) | 2.36(0.031) | -1.50(0.137) | -3.87(0.001) |
|  | R SMG | 17.44(<0.001) | 2.68(0.009) | -3.21(0.003) | -5.90(<0.001) |
|  | R PrCGsup | 15.26(<0.001) | 3.48(0.001) | -1.98(0.052) | -5.46(<0.001) |
|  | R SM | 4.60(0.013) | 2.00(0.074) | -0.97(0.335) | -2.98(0.012) |

**Supplementary Table 2.** One-way ANOVA and post hoc analysis of intrinsic functional connectivity. The F values and associated P values represent the main effect of the group from a Mixed-design ANOVA. The reported p values have been corrected for multiple comparisons across the ROIs using the FDR method. The t values and associated P values are from post hoc pairwise comparisons with FDR correction.

| Seed: **LSTG** | Target regions | Group main effect<br>$F_{2,69}(P_{fdr})$ | OMs vs. ONMs<br>$t_{69}(P_{fdr})$ | OMs vs. YNMs<br>$t_{69}(P_{fdr})$ | ONMs vs. YNMs<br>$t_{69}(P_{fdr})$ |
| --- | --- | --- | --- | --- | --- |
|  | L SMA | 6.20(0.010) | -0.66(0.511) | -3.52(0.002) | -2.11(0.058) |
|  | L SMG | 5.68(0.010) | -0.17(0.869) | -3.36(0.004) | -2.38(0.031) |
|  | L PrCGsup | 4.37(0.022) | -0.43(0.668) | -2.96(0.013) | -1.87(0.099) |
|  | L SM | 2.80(0.068) | -0.55(0.585) | -2.35(0.064) | -1.33(0.281) |
| Seed: **RSTG** | Target regions |  |  |  |  |
|  | R SMA | 5.23(0.015) | 0.05(0.964) | -3.20(0.006) | -2.42(0.027) |
|  | R SMG | 5.27(0.015) | -0.81(0.420) | -3.22(0.006) | -1.78(0.120) |
|  | R PrCGsup | 3.31(0.050) | 0.06(0.954) | -2.54(0.040) | -1.94(0.085) |
|  | R SM | 3.12(0.050) | -0.71(0.481) | -2.47(0.048) | -1.29(0.301) |

**Supplementary Table 3.**
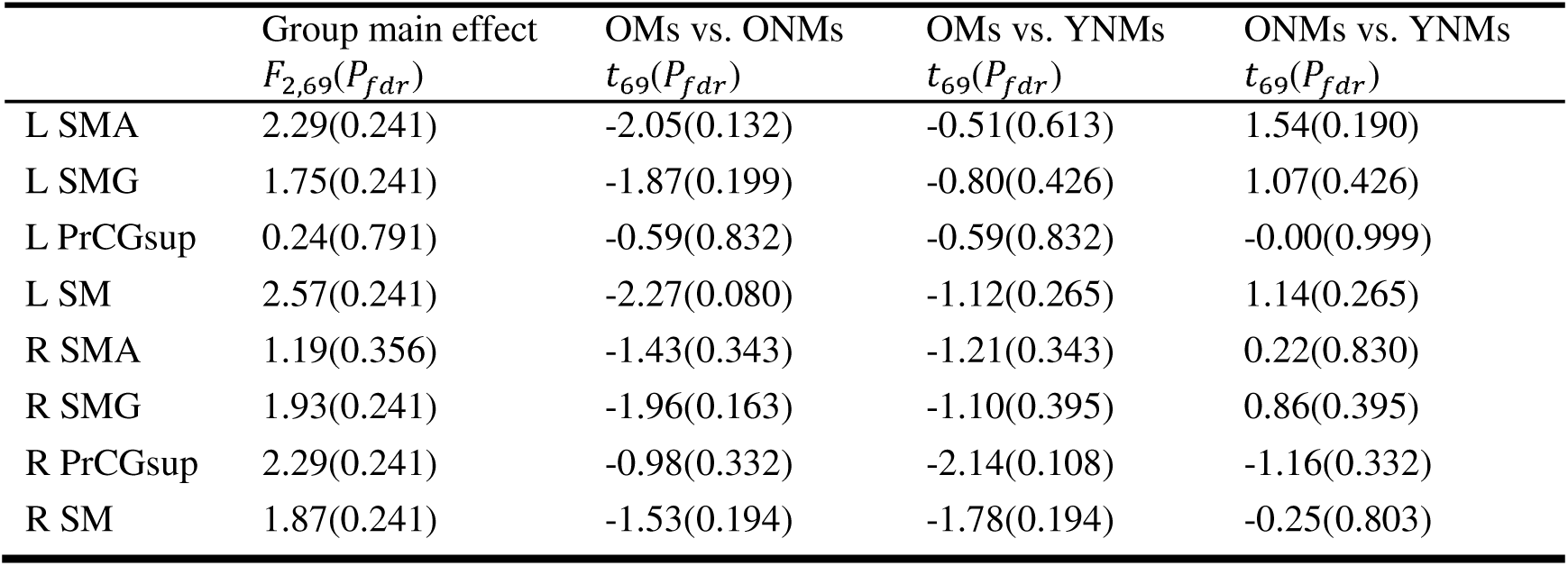
Mixed-design ANOVA and post hoc analysis of BOLD activation. The F values and associated P values represent the main effect of the group from a Mixed-design ANOVA. The reported P values have been corrected for multiple comparisons across the ROIs using the FDR method. The t values and associated P values are from post hoc pairwise comparisons with FDR correction.

## References

1. N. D. Anderson, F. I. M. Craik, 50 Years of Cognitive Aging Theory. The Journals of Gerontology: Series B 72, 1–6 (2017).

2. D. C. Park, P. Reuter-Lorenz, The Adaptive Brain: Aging and Neurocognitive Scaffolding. Annual Review of Psychology 60, 173–196 (2009).

3. D. C. Park, S. B. Festini, Theories of Memory and Aging: A Look at the Past and a Glimpse of the Future. The Journals of Gerontology: Series B 72, 82–90 (2017).

4. P. A. Reuter-Lorenz, D. C. Park, How Does it STAC Up? Revisiting the Scaffolding Theory of Aging and Cognition. Neuropsychol Rev 24, 355–370 (2014).

5. R. Cabeza, Hemispheric asymmetry reduction in older adults: The HAROLD model. Psychology and Aging 17, 85–100 (2002).

6. R. Cabeza, et al., Maintenance, reserve and compensation: the cognitive neuroscience of healthy ageing. Nat Rev Neurosci 19, 701–710 (2018).

7. Y. Stern, et al., Whitepaper: Defining and investigating cognitive reserve, brain reserve, and brain maintenance. Alzheimer’s & Dementia 16, 1305–1311 (2020).

8. S. W. Davis, N. A. Dennis, S. M. Daselaar, M. S. Fleck, R. Cabeza, Qué PASA? The Posterior–Anterior Shift in Aging. Cerebral Cortex 18, 1201–1209 (2008).

9. P. A. Reuter-Lorenz, K. A. Cappell, Neurocognitive Aging and the Compensation Hypothesis. Curr Dir Psychol Sci 17, 177–182 (2008).

10. Y. Stern, Cognitive Reserve and Alzheimer Disease. Alzheimer Disease & Associated Disorders 20, S69 (2006).

11. W. Kim, et al., An fMRI Study of Differences in Brain Activity Among Elite, Expert, and Novice Archers at the Moment of Optimal Aiming. Cognitive and Behavioral Neurology 27, 173 (2014).

12. G. Bernardi, et al., How Skill Expertise Shapes the Brain Functional Architecture: An fMRI Study of Visuo-Spatial and Motor Processing in Professional Racing-Car and Naïve Drivers. PLOS ONE 8, e77764 (2013).

13. Y. Du, R. J. Zatorre, Musical training sharpens and bonds ears and tongue to hear speech better. Proceedings of the National Academy of Sciences of the United States of America 114, 13579–13584 (2017).

14. A. Costa, N. Sebastián-Gallés, How does the bilingual experience sculpt the brain? Nature reviews. Neuroscience 15, 336 (2014).

15. E. B. J. Coffey, N. B. Mogilever, R. J. Zatorre, Speech-in-noise perception in musicians: A review. Hearing Research 352, 49–69 (2017).

16. R. J. Zatorre, J. L. Chen, V. B. Penhune, When the brain plays music: auditory-motor interactions in music perception and production. Nat Rev Neurosci 8, 547–558 (2007).

17. G. Schlaug, “Chapter 3 - Musicians and music making as a model for the study of brain plasticity” in *Progress in Brain Research*, Music, Neurology, and Neuroscience: Evolution, the Musical Brain, Medical Conditions, and Therapies., E. Altenmüller, S. Finger, F. Boller, Eds. (Elsevier, 2015), pp. 37–55.

18. S. C. Herholz, R. J. Zatorre, Musical Training as a Framework for Brain Plasticity: Behavior, Function, and Structure. Neuron 76, 486–502 (2012).

19. X. Jin, L. Zhang, G. Wu, X. Wang, Y. Du, Compensation or preservation? Different roles of functional lateralization in speech perception in older non-musicians and musicians. [Preprint] (2023). Available at: https://www.biorxiv.org/content/10.1101/2023.04.19.537446v1 [Accessed 10 April 2024].

20. L. Zhang, X. Wang, C. Alain, Y. Du, Successful aging of musicians: Preservation of sensorimotor regions aids audiovisual speech-in-noise perception. Science Advances 9, eadg7056 (2023).

21. S. Anderson, T. White-Schwoch, A. Parbery-Clark, N. Kraus, A dynamic auditory-cognitive system supports speech-in-noise perception in older adults. Hearing Research (2013). 10.1016/j.heares.2013.03.006.

22. E. B. J. Coffey, N. B. Mogilever, R. J. Zatorre, Speech-in-noise perception in musicians: A review. Hearing Research (2017). 10.1016/j.heares.2017.02.006.

23. D. Fleming, S. Belleville, I. Peretz, G. West, B. R. Zendel, The effects of short-term musical training on the neural processing of speech-in-noise in older adults. Brain and Cognition 136 (2019).

24. E. Dubinsky, E. A. Wood, G. Nespoli, F. A. Russo, Short-Term Choir Singing Supports Speech-in-Noise Perception and Neural Pitch Strength in Older Adults With Age-Related Hearing Loss. Frontiers in Neuroscience 13, 1153 (2019).

25. E. B. J. Coffey, A. M. P. Chepesiuk, S. C. Herholz, S. Baillet, R. J. Zatorre, Neural Correlates of Early Sound Encoding and their Relationship to Speech-in-Noise Perception. Frontiers in Neuroscience 11 (2017).

26. C. Alain, B. R. Zendel, S. Hutka, G. M. Bidelman, Turning down the noise: The benefit of musical training on the aging auditory brain. Hearing Research 308, 162–173 (2014).

27. B. L. Giordano, et al., Contributions of local speech encoding and functional connectivity to audio-visual speech perception. eLife 6, e24763 (2017).

28. A. Keitel, J. Gross, C. Kayser, Shared and modality-specific brain regions that mediate auditory and visual word comprehension. eLife 9, 1–23 (2020).

29. H. Park, R. A. A. Ince, P. G. Schyns, G. Thut, J. Gross, Representational interactions during audiovisual speech entrainment: Redundancy in left posterior superior temporal gyrus and synergy in left motor cortex. PLoS Biology 16, e2006558 (2018).

30. H. Park, C. Kayser, G. Thut, J. Gross, Lip movements entrain the observers’ low-frequency brain oscillations to facilitate speech intelligibility. eLife 5, e14521 (2016).

31. L. Zhang, Y. Du, Lip movements enhance speech representations and effective connectivity in auditory dorsal stream. Neuroimage 257, 119311 (2022).

32. Y. Du, B. R. Buchsbaum, C. L. Grady, C. Alain, Increased activity in frontal motor cortex compensates impaired speech perception in older adults. Nature Communications 7, 1–12 (2016).

33. P. C. M. Wong, et al., Aging and cortical mechanisms of speech perception in noise. Neuropsychologia 47, 693–703 (2009).

34. I. Wollman, V. Penhune, M. Segado, T. Carpentier, R. J. Zatorre, Neural network retuning and neural predictors of learning success associated with cello training. Proceedings of the National Academy of Sciences 115, E6056–E6064 (2018).

35. G. Hickok, D. Poeppel, The cortical organization of speech processing. Nature Reviews Neuroscience 8, 393–402 (2007).

36. J. P. Rauschecker, Where, When, and How: Are they all sensorimotor? Towards a unified view of the dorsal pathway in vision and audition. Cortex 98, 262–268 (2018).

37. C. Alain, S. R. Arnott, S. Hevenor, S. Graham, C. L. Grady, “What” and “where” in the human auditory system. Proc. Natl. Acad. Sci. U.S.A. 98, 12301–12306 (2001).

38. L. E. Bernstein, E. Liebenthal, Neural pathways for visual speech perception. Frontiers in Neuroscience 8, 1–18 (2014).

39. L. Zhang, X. Fu, D. Luo, L. Xing, Y. Du, Musical Experience Offsets Age-Related Decline in Understanding Speech-in-Noise: Type of Training Does Not Matter, Working Memory Is the Key. Ear and Hearing 42, 258–270 (2021).

40. H. A. Manan, E. A. Franz, A. N. Yusoff, S. Z. M. S. Mukari, The effects of aging on the brain activation pattern during a speech perception task: an fMRI study. Aging Clinical and Experimental Research 27, 27–36 (2015).

41. P. Tremblay, V. Brisson, I. Deschamps, Brain aging and speech perception: Effects of background noise and talker variability. NeuroImage 227, 117675 (2021).

42. P. Tremblay, A. S. Dick, S. L. Small, Functional and structural aging of the speech sensorimotor neural system: functional magnetic resonance imaging evidence. Neurobiology of Aging 34, 1935–1951 (2013).

43. M. Bilodeau-Mercure, C. L. Lortie, M. Sato, M. J. Guitton, P. Tremblay, The neurobiology of speech perception decline in aging. Brain Struct Funct 220, 979–997 (2015).

44. J. V. Strong, B. T. Mast, The cognitive functioning of older adult instrumental musicians and non-musicians. *Aging*, Neuropsychology, and Cognition 26, 367–386 (2019).

45. C. Giacosa, F. J. Karpati, N. E. V. Foster, V. B. Penhune, K. L. Hyde, Dance and music training have different effects on white matter diffusivity in sensorimotor pathways. NeuroImage 135, 273–286 (2016).

46. G. F. Halwani, P. Loui, T. Rüber, G. Schlaug, Effects of Practice and Experience on the Arcuate Fasciculus: Comparing Singers, Instrumentalists, and Non-Musicians. Front. Psychology 2 (2011).

47. X. Li, R. J. Zatorre, Y. Du, The Microstructural Plasticity of the Arcuate Fasciculus Undergirds Improved Speech in Noise Perception in Musicians. Cerebral Cortex 31, 3975–3985 (2021).

48. Y. Du, B. R. Buchsbaum, C. L. Grady, C. Alain, Noise differentially impacts phoneme representations in the auditory and speech motor systems. Proceedings of the National Academy of Sciences of the United States of America 111, 7126–31 (2014).

49. S. L. Mattys, M. H. Davis, A. R. Bradlow, S. K. Scott, Speech recognition in adverse conditions: A review. Language and Cognitive Processes 27, 953–978 (2012).

50. B. Fauvel, et al., Morphological brain plasticity induced by musical expertise is accompanied by modulation of functional connectivity at rest. NeuroImage 90, 179–188 (2014).

51. X. Zhang, P. Tremblay, Aging of Amateur Singers and Non-singers: From Behavior to Resting-state Connectivity. Journal of Cognitive Neuroscience 35, 2049–2066 (2023).

52. M.-Á. Palomar-García, R. J. Zatorre, N. Ventura-Campos, E. Bueichekú, C. Ávila, Modulation of Functional Connectivity in Auditory–Motor Networks in Musicians Compared with Nonmusicians. Cerebral Cortex 27, 2768–2778 (2017).

53. F. Liem, L. Geerligs, J. S. Damoiseaux, D. S. Margulies, “Functional connectivity in aging” in *Handbook of the Psychology of Aging*, (Elsevier, 2021), pp. 37–51.

54. L. K. Ferreira, G. F. Busatto, Resting-state functional connectivity in normal brain aging. Neuroscience & Biobehavioral Reviews 37, 384–400 (2013).

55. J. Yu, J. Li, X. Huang, The Beijing version of the montreal cognitive assessment as a brief screening tool for mild cognitive impairment: a community-based study. BMC Psychiatry 12, 156 (2012).

56. H.-W.-S. Bao, bruceR: BRoadly Useful Collections and Extensions of R functions. (2020).

57. R. W. Cox, AFNI: Software for analysis and visualization of functional magnetic resonance neuroimages. Computers and Biomedical Research 29, 162–173 (1996).

58. D. G. McLaren, M. L. Ries, G. Xu, S. C. Johnson, A generalized form of context-dependent psychophysiological interactions (gPPI): A comparison to standard approaches. NeuroImage 61, 1277–1286 (2012).

59. J. Wang, et al., GRETNA: a graph theoretical network analysis toolbox for imaging connectomics. Frontiers in Human Neuroscience 9 (2015).

60. Y. Tong, et al., Seeking Optimal Region-Of-Interest (ROI) Single-Value Summary Measures for fMRI Studies in Imaging Genetics. PLOS ONE 11, e0151391 (2016).

61. M. Koller, robustlmm: an R package for robust estimation of linear mixed-effects models. Journal of statistical software 75, 1–24 (2016).

